# MedMCP: An Open Agentic Ecosystem for Medical Imaging Workflows

**DOI:** 10.64898/2026.10.12.758708

**Authors:** Julian McGinnis, Paul Friedrich, Mark Mühlau, Jan S. Kirschke, Martin Styner, Daniel Rueckert, Philippe C. Cattin, Benedikt Wiestler

**Author notes:** Shared first authorship (Julian McGinnis, Paul Friedrich).

## Abstract

Deep learning has driven rapid progress in medical image analysis, with methods for tasks such as organ segmentation, anomaly detection, and radiology report generation now matching or surpassing expert performance on several benchmarks. While many methods are released as open source, their routine adoption by clinicians and medical researchers remains limited. The bottleneck is not a lack of capable tools, but the absence of infrastructure to make them accessible to non-technical users.

To break this barrier, we present **MedMCP**, an agentic framework that solves this *last-mile problem* along two axes: (1) a *local agentic workspace* that lets users apply state-of-the-art image analysis methods through natural language, and (2) a *community-driven tool ecosystem* that lets developers deploy and distribute capabilities as validated, independently testable servers.

MedMCP enforces a strict orchestration-execution boundary in which a local language model translates user intent into workflow plans, while all image analysis is delegated to domain-specific tools. The framework runs entirely on-premise, every side-effecting operation passes through a human-in-the-loop permission gate, and a full provenance trail links each output to the tools, parameters, and inputs that produced it. An inspectable workspace lets users verify intermediate results rather than blindly trusting the agent, and completed sessions distill into deterministic, shareable workflows that re-execute without the agent, turning one-off interactions into reproducible analyses.

This paper introduces the project’s general idea and describes the design principles that guide its development. Beyond a single toolkit, we envision MedMCP as a community-driven platform where each contributed tool becomes easily accessible, compounding individual efforts into shared capability.

https://medmcp.ai

https://github.com/medmcp

## 1 Introduction

The medical imaging community has produced a rich ecosystem of well-established, open-source tools for diverse tasks ranging from brain extraction (Isensee et al., 2019) and multi-modal registration (Tustison et al., 2021) to deep learning-based segmentation (Isensee et al., 2021; Wasserthal et al., 2023), computational pathology (Lu et al., 2021), or radiology report generation (Tanno et al., 2025). Yet a fundamental problem persists: most practitioners who stand to benefit from these tools cannot use them (Kelly et al., 2019; Cabitza et al., 2020). We argue that, beyond regulatory hurdles (Muehlematter et al., 2021), this barrier is not due to inadequate tools but rather to the level of technical fluency required to operate them. Clinicians rarely possess this level of technical fluency, nor should they need to.

We therefore present MedMCP, an agentic ecosystem that aims to close this gap by exposing medical imaging capabilities through natural language. Its central design principle is a strict separation between orchestration and execution: a local large language model (LLM) interprets user intent, sequences operations, and invokes validated tools, while all image processing is delegated to those tools. This separation is what makes the framework both trustworthy and extensible. The LLM never touches the data directly, and new capabilities can be added without retraining or modifying the orchestrator. The entire system runs on-premise, so data never leaves the institution, satisfying the governance requirements of clinical and translational research environments.

Natural-language access alone, however, is of little value if users cannot verify and trust what the system produces. We therefore build MedMCP into a complete agentic workspace rather than a bare conversational interface. It pairs the chat with a file explorer and image viewer for inspecting inputs and intermediate results, a full provenance trail linking every output to the tools, parameters, and inputs that produced it, and a human-in-the-loop permission gate through which each side-effecting operation, i.e. every operation that interacts with data, must pass. Completed sessions can be distilled into deterministic workflows that re-execute without the agent and can be shared with others, turning one-off interactions into reproducible, reusable analyses.

The remainder of this paper further establishes the problem that motivates such a framework (Section 2), describes the design philosophy behind its architectural decisions (Section 3), details the system’s (current) implementation (Section 4), and discusses its potential impact. This paper is meant as a **“living document”**, i.e., we will regularly update it to keep track with MedMCP’s developments and will add concrete use cases, systematic performance reports, and experience from real-world deployments.

## 2 Motivation

### 2.1 The Last-Mile Problem in Medical Imaging

The motivation for this project stems from a simple observation:

**The Last-Mile Problem**. Despite a mature ecosystem of medical imaging tools, the practitioners best positioned to apply them are systematically unable to do so. The barrier is not capability, it is the accumulated technical complexity required to operate these tools. We refer to this persistent disconnect between **tool availability** and **tool accessibility** as the **Last-Mile Problem**.

We argue that this gap persists because of a structural mismatch between the involved communities:

1. On the **developer side**, researchers who produce methods operate within an incentive structure that rewards publication, not adoption. For most methods, the *de facto* accessibility standard is a public code repository with an explanatory README file (Moassefi et al., 2023; Simkó et al., 2024). Anything beyond this threshold is left to the initiative of researchers who aim to sustain their work beyond the publication. Packaging a tool for non-technical users is a distinct engineering effort, encompassing installation testing, multi-platform support, error handling for unexpected inputs, and ongoing maintenance, none of which is rewarded by academic metrics. Projects such as FreeSurfer (Fischl, 2012), nnU-Net (Isensee et al., 2021), and TotalSegmentator (Wasserthal et al., 2023) represent the upper bound of what sustained accessibility investment currently achieves, offering extensive documentation, containerization, and stable interfaces. Yet even these tools are rarely used by clinicians. Processing a cohort with FreeSurfer remains a notable programming effort for domain researchers, and nnU-Net models are, in practice, trained and deployed by machine learning researchers rather than radiologists.
2. On the **user side**, applying even a single state-of-the-art imaging pipeline requires fluency in command-line interfaces, environment management, file format conventions, and tool-specific APIs, i.e., expertise accumulated over years of research-first development that domain specialists neither have nor should be expected to acquire (Gorgolewski et al., 2017). Faced with this overhead, clinical and imaging research users default to established workflows and forgo the capabilities of newer methods. Beyond the impact on tool choices, the overhead constrains research output itself: studies devised by domain experts may never be conducted or become dependent on the availability of technical collaborators, thereby gating clinical research questions behind engineering capacity. The limiting factor is not unwillingness to adopt better tools, but the cost of integrating them.

The result is a persistent, structural barrier between working methods and unreachable users. *This observation is not new*. The medical imaging community has long recognized that accessibility is a first-class challenge in translational research. Individual efforts to address the problem through tutorials, containerization, or web interfaces have, however, remained fragmented and method-specific (Kelly et al., 2019; Von Chamier et al., 2021; Varoquaux & Cheplygina, 2022).

#### We envision and develop MedMCP as a community-driven effort to address this challenge systematically

Rather than placing the accessibility burden on individual developers, the framework provides shared, easy-to-use infrastructure. The core components are maintained centrally, so contributors supply the domain capability only and inherit natural-language access, permission gating, provenance tracking and the deterministic replay engine with shareable workflows without additional effort. Each contribution becomes available to users of the framework and tool accessibility scales with the community.

### 2.2 Why Not Prompt a General-Purpose LLM?

While prompting a general-purpose LLM may seem superficially attractive, it is deeply unreliable (Bhayana et al., 2023; Bhayana, 2024), and closer inspection reveals several fundamental limitations that render this approach unsuitable for medical imaging workflows.

The most basic problem is unreliable execution. Dynamic code generation may succeed intermittently, but it offers no guarantees (Liu et al., 2023; Tian et al., 2025). Small variations in input format, library versions, or task phrasing can produce silently incorrect results or outright failures (Sclar et al., 2024; Ouyang et al., 2025). Additionally, executing a workflow through generated code requires the model to reproduce functionality that is already robustly implemented, tested, and validated in established libraries. Regenerating this logic from scratch discards years of accumulated engineering effort and introduces the risk of subtle deviations from the very reference implementations whose correctness the community already trusts.

The model’s susceptibility to hallucination amplifies the risks. LLMs readily invent non-existent function signatures, parameters or file conventions (Patil et al., 2024; Spracklen et al., 2025; Zhang et al., 2025). Without a fixed, validated tool interface to constrain them, these hallucinations manifest as runtime errors at best and incorrect but plausible-looking outputs at worst. Free-form code execution is inherently opaque, and this makes any clean fix difficult. There’s no natural checkpoint at which to gate permissions for side-effecting operations, no provenance trail tying outputs back to the inputs and parameters that produced them, and no way for a non-technical user to see what the generated code actually did.

Finally, even when code is generated correctly, the user remains responsible for providing a working execution environment with the correct dependencies, versions, and system configuration, reintroducing exactly the technical burden the framework is meant to eliminate (Pimentel et al., 2019; Trisovic et al., 2022).

MedMCP avoids these failure modes by construction. The orchestrator never generates code that touches the data but instead selects among a fixed set of validated tools with well-defined interfaces, an approach shown to improve reliability and reduce API hallucination relative to free-form generation (Schick et al., 2023; Yao et al., 2023; Patil et al., 2024; Qin et al., 2024). This constrains the model to a space of operations known to be correct, makes every action observable and gateable, and shifts the environmental burden from the user to the tool packager, who resolves it once on behalf of the community.

## 3 Design Philosophy

MedMCP is built upon four core design principles that we consider essential for a medical agent to succeed:

1. **Accessibility First**. MedMCP exists to make validated imaging tools reachable, not to produce new ones. Every architectural decision is evaluated against one question: *Does this reduce the barrier between a working method and the practitioner who needs it?*
2. **Bounded by Design**. The agent’s role is strictly limited to invoking pre-validated tools. It is not meant to write code or improvise solutions. The value of the framework lies in the reliability of known, tested behavior, not in creative problem-solving.
3. **Local Execution Only**. Medical data is subject to strict privacy regulations and institutional governance policies. The system is therefore required to run entirely on-premise. No imaging data, patient meta-data, or intermediate results leave the institution’s infrastructure. This is a hard architectural constraint, not an optional feature.
4. **Community-Driven Growth**. No single team can cover the full breadth of medical imaging workflows across radiology, pathology, cardiology, and beyond. MedMCP is designed to grow through community contributions: a shared schema for tool and skill meta-data, and a central registry for discovery ensure that each new contribution is immediately available to all users.

## 4 Implementation

MedMCP consists of three main components: (1) a *workspace* in which the user operates, (2) an *orchestrator* that translates requests into executable plans, and (3) a set of *tool stacks* that carry out the computation. These components communicate through open protocols and a shared data directory (Figure 1). All components are distributed as multi-architecture container images and run on hardware ranging from single GPU-enabled workstations to servers and clusters.

**Figure 1.**
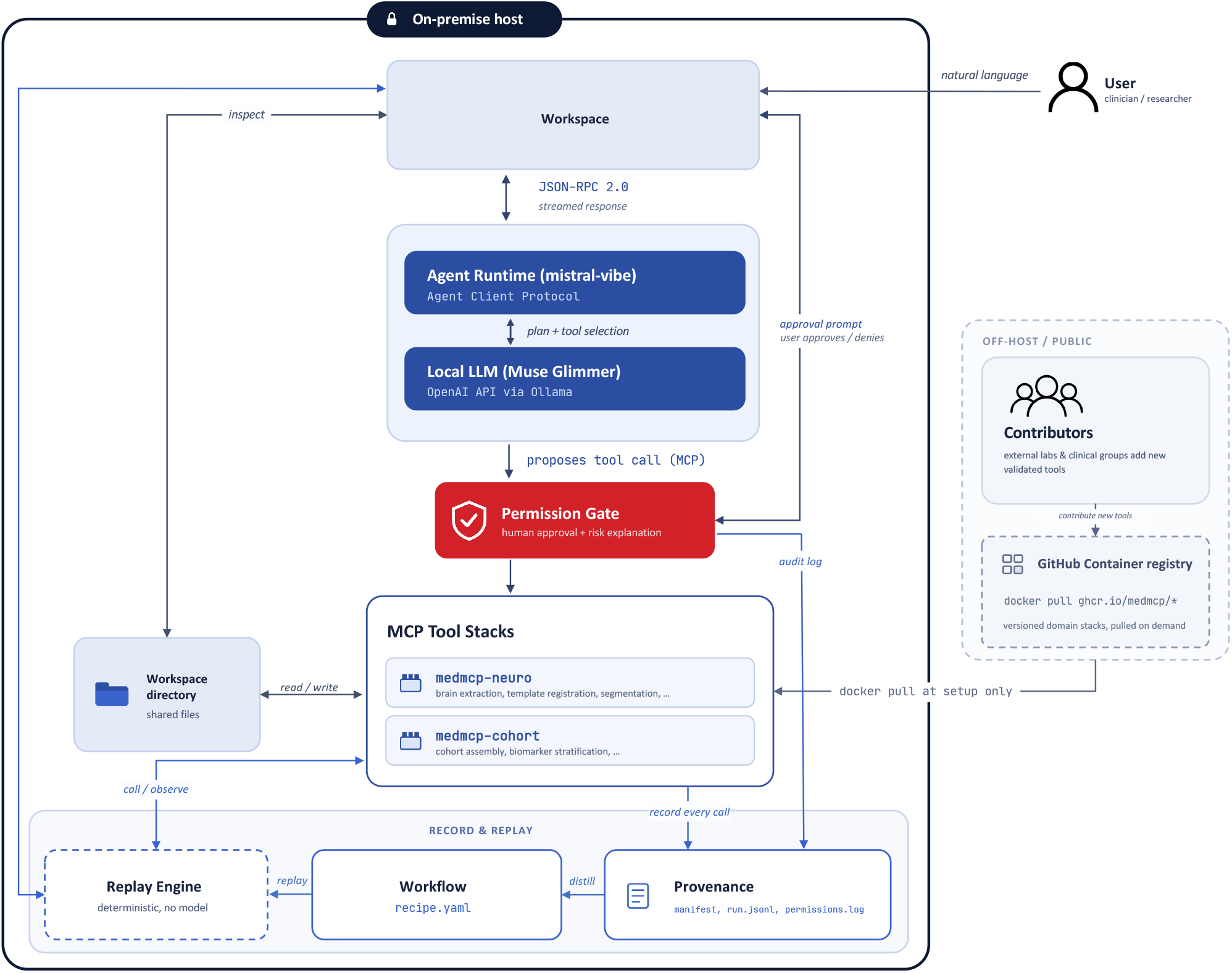
An overview of the MedMCP architecture. The *user* interacts with the *workspace* via natural language, accesses the MCP stacks’ capabilities, observes the workspace directory and interacts with the replay engine from there. Imaging capabilities are grouped in separate *tool stacks* that are community-contributed MCP servers gated by a human-in-the-loop permission gate. Every interaction, permission, and tool call, is logged and can be distilled into a *workflow* that can be replayed without the agent in the loop.

The workspace controls the orchestrator via the Agent Client Protocol (ACP)^1^, a JSON-RPC 2.0 interface carried over the orchestrator’s standard input and output. The orchestrator, in turn, accesses all tools through the Model Context Protocol (MCP)^2^, which is becoming the *de facto* standard for connecting language models to external tools. The model itself is served locally behind an OpenAI-compatible endpoint^3^. As all interfaces are public and implementation-independent, each layer can be replaced in isolation: a newer model, a different inference server, or tools written in other languages can be integrated without modifying MedMCP.

### 4.1 One Request, End to End

We introduce MedMCP’s architecture by tracing an exemplary interaction through the entire system. Consider a user who opens a T1-weighted scan in the workspace viewer and provides the following instruction: *“remove the skull from this scan and register it to MNI space”*.

Originating from the UI, the workspace (Section 4.2) sends the request to the orchestrator along with a note identifying the currently displayed file, so that “this scan” resolves to the user’s currently selected file rather than requiring the model to guess. The orchestrator (Section 4.3) appends the request to the conversation and instructs the local model to continue, supplying the descriptions of every tool that the installed stacks (Section 4.4) currently expose. The model answers with either text or a structured tool call, i.e., the tool name and its arguments.

Proposed tool calls are not just executed, but are gated by approval of the user (Section 4.5), who is provided with a dialog listing the tool, tool arguments and intended usage, accompanied by a one-line, plain-language summary of the workflow and its associated risk level from a predefined risk label set indicating permission-levels. The call is only forwarded, *via MCP*, to the stack that owns the required tool after user approval. The stack executes the validated implementation, in this case brain extraction and registration, reading and writing files in the shared workspace directory. The file explorer and viewer periodically refresh, so that a newly appearing brain mask is rendered visible. The tool call is appended to the session’s provenance record (Section 4.6). The result is forwarded to the conversation, and the model/user loop is repeated until the model produces the final textual answer to the user’s instruction. By design, the complete exchange remains local on the host. No image, identifier, or intermediate result leaves the local setup.

### 4.2 Workspace

The workspace is implemented as a FastAPI^4^ server that serves a React and TypeScript application. In the current version, we provide a four-panel design with a file explorer, a medical-image viewer, a replay engine, and the agent chat. An example view is shown in Figure 2.

**Figure 2.**
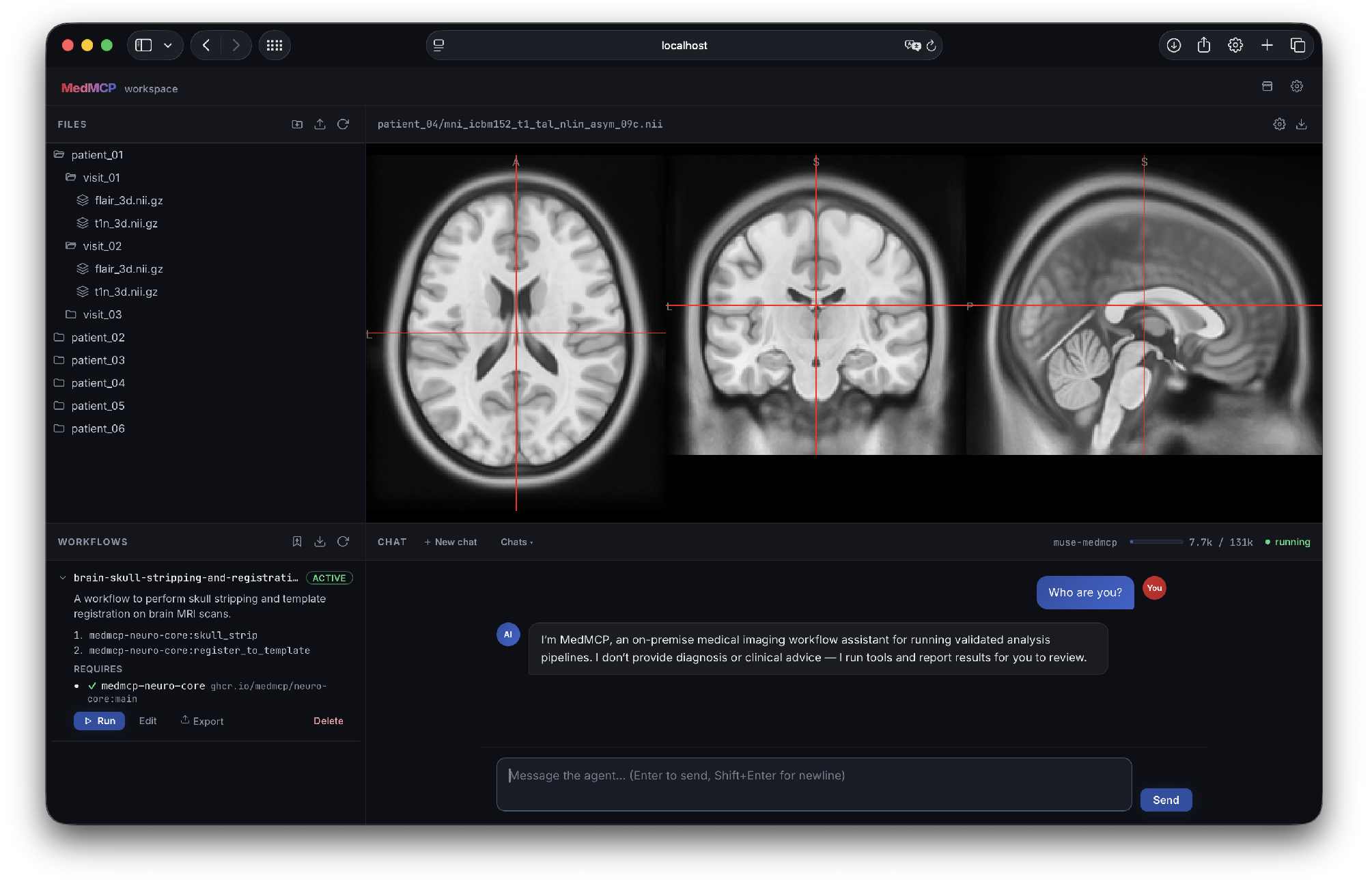
An example view of the MedMCP workspace containing an interactive file explorer *(top left)*, a medical image viewer with support for all common file formats *(top right)*, a replay engine for (re-)running distilled workflows *(bottom left)*, and the chat interface *(bottom right)* that is used for calling tools and handling permission gating.

MedMCP interacts with a single directory only. This directory serves as the root of the file explorer, the source for the viewer, and the working directory for the agent’s tools, so the user always sees the files the tools read and produce. The viewer selects a renderer by file type, i.e. volumetric images are shown in an interactive multiplanar and 3D display, built with Niivue^5^, that also supports overlaying segmentation masks over its source image, while documents, code, 2D images, and tables fall back to appropriate previews. Explorer and viewer reload whenever a tool call completes, or a turn ends, so results appear without a manual refresh.

The chat panel streams the model’s answer as it is generated, renders each tool call as an inspectable card, hosts the approval dialogs, and reports the model in use and how much of its context window is occupied. The workspace is also used to interact with persistent conversations.

### 4.3 Orchestration

The orchestrator is a long-running agent-runtime process that owns the conversation loop. One process serves every open chat, with sessions kept apart by identifier. Our local LLM inference is currently handled by Ollama^6^, serving an open-weights Muse Glimmer model ^7^ for our reference implementation. We also provide an easy way to change the model depending on the available hardware and will regularly update our reference implementation.

The model’s job is narrow by construction. On each turn, it emits either text for the user or the name of a tool with its arguments. It never emits code that is executed against the data, and it never sees the data itself, as images are passed between tools as file paths. Two mechanisms steer its behavior without any retraining: (1) a *system prompt* that fixes the scope of the assistant, and (2) *skills* that describe how a stacks tools work and how they are invoked correctly.

### 4.4 Tool Stacks

A *tool stack* is a unit of extension, i.e., one MCP server exposing a related family of tools, packaged with its own isolated environment, its own pinned dependencies, and its own skills. Isolation is what makes independent contribution practical, as two stacks may require incompatible versions of the same framework without either affecting the other or the core.

Installing a new stack is a single action in the workspace. The user picks it from a catalog of available stacks, MedMCP pulls the docker image, reads the label, extracts the skills, and records a small launch manifest. From that point, the stacks’ tools become available across all sessions.

### 4.5 Permission Gate

Every tool call proposed by the model is intercepted before execution and released only upon an explicit user decision. There is no auto-approval path in the system. An example permission gate is shown in Figure 3.

**Figure 3.**
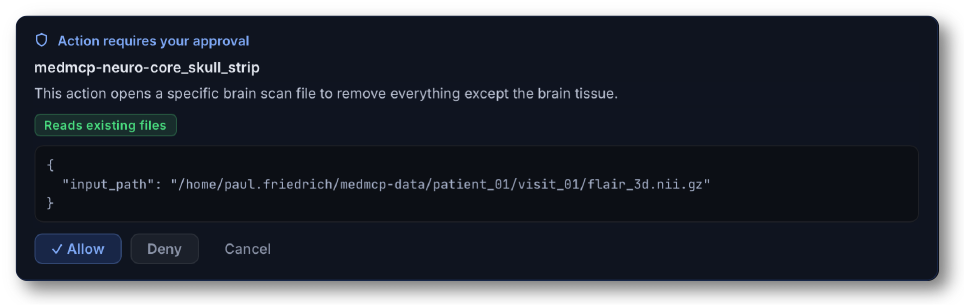
An example permission gate for the *skull stripping* tool. The user sees the tool name, a natural language explanation, and the arguments used for invoking the tool (in this case only the input path).

The dialog shows the tool, the arguments it would be called with, and a plain-language sentence explaining what the action will do and what will change as a result. This sentence is produced by a short, separate pass through the same LLM, prompted to write for a reader with no technical background and to tag the call against a fixed taxonomy of risk categories. Every decision, approval, or rejection is written to an append-only audit log.

This gate exists as the LLM can be steered by the content it processes, e.g., a referral letter, a report, or a tool’s own output may contain text that reads as an instruction. Rather than attempting to detect such attempts, the architecture requires that a person authorizes each side-effecting action.

### 4.6 Provenance

Each conversation accumulates an append-only record. A manifest, written with the first request, captures the execution environment, i.e., the framework revision, the installed stacks with their versions or image digests, the active model and its parameters, and the platform. One normalized entry per completed tool call is then appended to a log, recording the stack, the tool, the arguments as they were finally resolved, the structured result, the user’s permission decision, the duration, and the outcome. The permission audit trail is mirrored alongside. From these files, a human-readable report can be rendered on demand that links every output back to the tool, parameters, and inputs that produced it.

### 4.7 Distilled Workflows and Deterministic Replay

A session that successfully produced results can be turned into a repeatable procedure. Distillation reads the session’s record, discards the calls that carry no reusable meaning, like exploration, failures, and anything the user rejected, and generalizes remaining tool calls by replacing the concrete file paths with named placeholders, separating the inputs the workflow consumes from the values earlier steps produce. The result is a pair of files: a machine-readable recipe and a human-readable description drafted by the local model, with a mechanical rendering as a fallback so distillation never fails outright. A workflow can be exported as a single self-contained file (steps, inputs, documentation, and the stacks it requires, pinned by version or image digest) and imported by other users, researchers or collaborators. An example workflow is shown in Figure 4.

**Figure 4.**
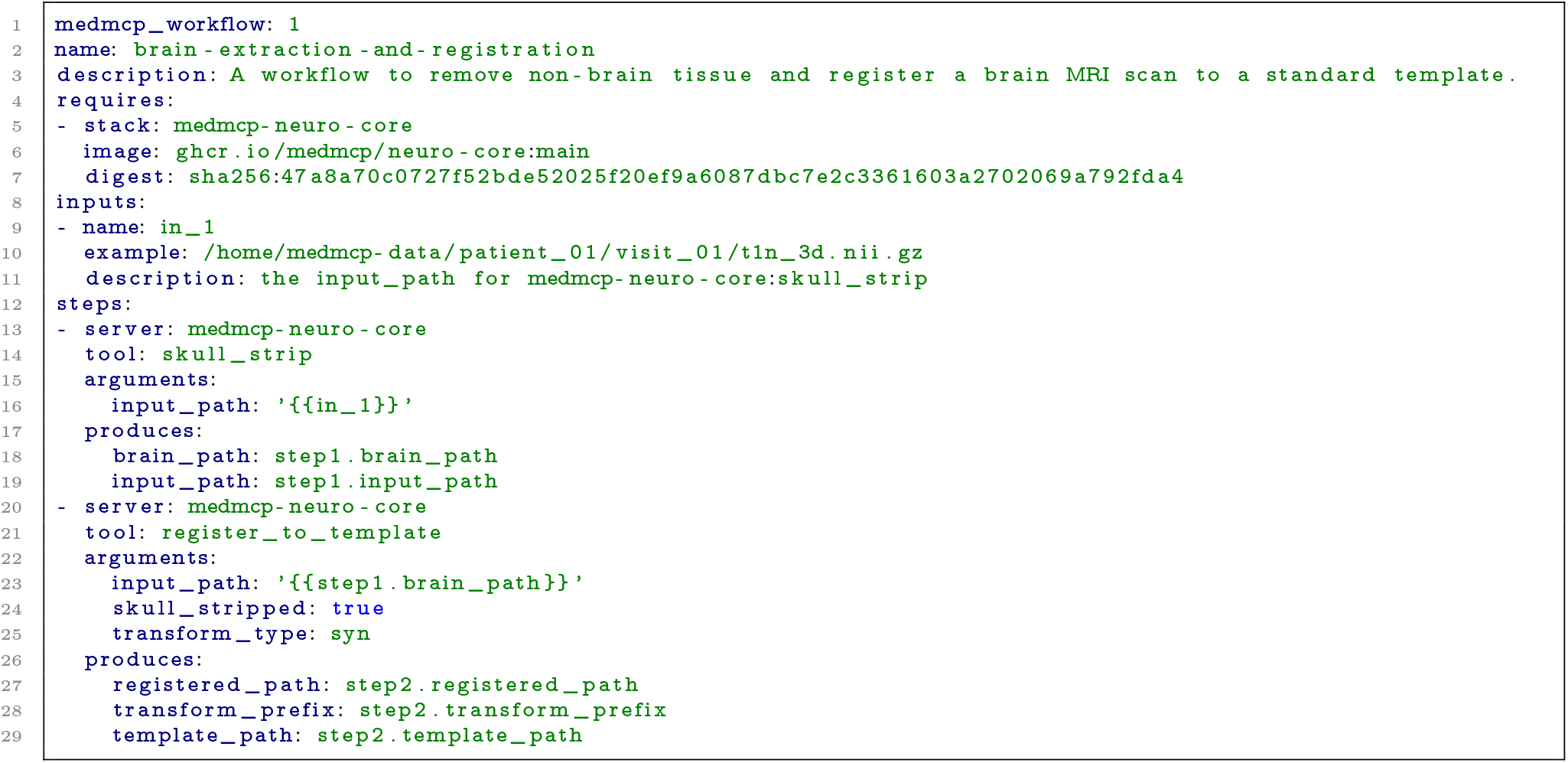
The workflow `brain-extraction-and-registration` that has been distilled from an example session in which the user first skull-stripped a T1 scan and then registered it to a template. The workflow can now be applied to another T1 scan.

A recipe re-executes through a replay engine that runs *without* the language model. The engine first validates the recipe, reporting missing inputs or stacks that are not installed, then substitutes the new inputs, starts the required tool servers, and executes the recorded steps in order, feeding each step’s declared outputs into the next step’s arguments exactly as in the original session. The same engine runs a whole cohort in one batch: the servers are started once and reused across all cases, each case is validated on its own inputs, and a failure in one does not abort the rest. Because replay deliberately bypasses both the model and the interactive permission gate, it is never triggered silently. The workspace shows the fully resolved steps and requires an explicit confirmation before the first tool runs.

Together, these three mechanisms determine what a session leaves behind. The permission gate governs what happens while it runs, provenance records what happened, and distillation converts what happened into something that can be run again, by someone else, on new data, without the model in the loop.

## 5 Call to Action

The argument of this paper is that accessibility scales with the community rather than a single team, which makes the following requests substantive rather than rhetorical.

### To method developers: *ship a stack alongside your paper*

A stack is one MCP server, its pinned environment, and its skills. Writing one is a bounded engineering task, considerably smaller than building a user-facing interface, and it is paid once. In exchange, a method becomes invocable in natural language, gated, logged, and replayable by users who would otherwise never run it. *We call for a stack being considered a normal release artifact of a methods paper, next to the repository and the model weights*. We provide easily adaptable tool stacks and containerization, CI/CD utilities and offer to host validated tools within the medmcp organization^8^.

### To clinicians and clinical researchers: *use it and tell us where it fails*

The failure modes that matter most are the ones encountered in real sessions with real data and real questions. Reports of failures such as incorrect tool selection, unclear approval dialogs, and results that appeared plausible but were not are valuable contributions. *We invite you to share your reports and desired functionality, including extensions, with us*, and to collaborate on resolving and integrating them.

## 6 Discussion

This paper accompanies the initial public release of MedMCP and describes a starting point and vision, not a finished system. MedMCP provides a general framework for making validated medical imaging tools usable through natural language, with local execution, human oversight and full provenance built in by design. Its defining choice is to constrain the agent to a fixed set of validated tools rather than allowing it to generate code at runtime. This restriction trades flexibility for reliability: requests without a matching tool cannot be served, but every operation that does run has known, tested behavior, and errors surface as visible failures rather than plausible-looking results (Dratsch et al., 2023; Patil et al., 2024). In a clinical research context, we consider this the appropriate balance. The practical consequence is that the utility of the framework scales with the coverage of its tool ecosystem, which is why MedMCP is designed around community contribution rather than centralized development.

Several limitations and open questions remain. Orchestration quality depends on the locally served language model, which may select an inappropriate tool or resolve arguments incorrectly. The approval dialog exposes each proposed call before execution, but this safeguard is only effective if users engage with it, and how well the permission gate holds up against approval fatigue in daily use is something only real deployments can tell us. Community-contributed stacks shift the burden of correctness to their contributors. The ecosystem will therefore require curation mechanisms, including registry review, standardized tests on reference data and machine-readable declarations of each tool’s validated domain, although which of these mechanisms work at scale is yet to be established. Finally, local inference presupposes local GPU resources, which constrains adoption in low-resourced settings.

The framework readily extends beyond its current scope. The initial stacks emphasize neuroimaging, reflecting the authors’ backgrounds, but the same packaging conventions apply to other imaging domains. Integration with institutional archives and community standards such as BIDS (Gorgolewski et al., 2017) will support cohort-scale studies, for which the batch replay engine already provides the execution substrate.

We are aware that we do not have all the answers yet. Which workflows practitioners actually attempt, where the orchestration fails, and what makes an approval dialog trustworthy rather than merely reassuring are empirical questions that this release is designed to surface, and this living document will evolve with the answers, incorporating user studies, orchestration benchmarks and experience from institutional deployments as they accumulate. None of the directions outlined here are settled, and we consider that a feature of the project rather than a shortcoming: **where MedMCP goes next should be shaped by the people who use and extend it**. We therefore explicitly invite discussion, whether on tool curation, orchestration models, clinical deployment, or directions we have not considered, and we are happy to engage. What we do propose with confidence is a change in how imaging methods are released: the stack as a standard companion of a methods paper, alongside the code repository and model weights. The contribution required of a developer is deliberately small, and in return a method becomes accessible to practitioners who would otherwise never run it. We anticipate that as the ecosystem grows and this journey unfolds alongside its community, MedMCP will help close the persistent gap between the pace of methodological progress in medical imaging and its adoption in clinical research practice.

## Acknowledgements

We would like to thank Florentin Bieder for helpful discussions around the project. This work is supported by the Munich Center for Machine Learning.

## Impact Statement

MedMCP removes the barrier between validated medical imaging methods and the practitioners who need them. Centers without dedicated imaging engineers gain access to state-of-the-art analysis, every result is traceable and replayable, and no data ever leaves the institution. Once *stacks* become standard companions of methods papers, each published method turns into a capability the entire community can use the day it is released. We are convinced that agentic frameworks of this kind will define how medical imaging research is conducted, with natural language as the interface and validated tools as the substrate, and with MedMCP we intend to actively shape that future rather than wait for it.

## Risk Assessment

The same lowered barrier that drives these benefits, however, also creates the risks we assess below. MedMCP is research software, not a medical device. It carries no regulatory clearance, and wrapping a tool in a conversational interface does not extend that tool’s intended use, its approved indication or the population in which it was validated. Outputs must not inform diagnostic or treatment decisions without independent verification by a qualified professional, and responsibility for regulatory compliance remains with the deploying institution.

The most serious risk the framework introduces is out-of-distribution use. The same interface that lets a clinician run a segmentation model without commandline experience also lets them run it on data the model was never validated for: a different scanner, sequence, field strength, age group or pathology. A manual invocation exposes at least some of these assumptions through documentation and parameters, while natural language hides them. Skills let tool authors declare intended use, and the workspace is built so that every intermediate result can be inspected rather than assumed. These mitigations are partial. A convention by which tools declare their validated domain in machine-readable form, and by which the orchestrator checks it against the data at hand and surfaces mismatches to the user, is in our view a necessary direction for the ecosystem.

A second risk is automation bias. The permission gate protects users only if they actually engage with it: fluent, plain-language explanations may themselves increase unwarranted trust, and repeated approval of routine calls tends toward reflexive confirmation (Dratsch et al., 2023). We deliberately provide no auto-approval path and no standing per-tool approvals, but we do not claim this eliminates the problem. Provenance and the inspectable workspace are intended to support verification, not to substitute for it.

Finally, local inference requires local hardware. A framework whose stated aim is democratization is nonetheless bounded by the availability of GPUs and could, in principle, widen rather than narrow the gap between well-resourced and low-resourced settings. Keeping the reference implementation runnable on modest, single-GPU workstations is therefore a design constraint we intend to maintain as the framework grows.

## Footnotes

1 https://agentclientprotocol.com

2 https://modelcontextprotocol.io

3 https://developers.openai.com/api/reference

4 https://fastapi.tiangolo.com

5 https://niivue.com

6 https://ollama.com

7 https://developer.meta.com/ai/models/muse-glimmer/

8 https://github.com/medmcp

## References

Bhayana, R. Chatbots and large language models in radiology: a practical primer for clinical and research applications. Radiology, 310(1):e232756, 2024.

Bhayana, R., Krishna, S., and Bleakney, R. R. Performance of chatgpt on a radiology board-style examination: insights into current strengths and limitations. Radiology, 307(5):e230582, 2023.

Cabitza, F., Campagner, A., and Balsano, C. Bridging the “last mile” gap between ai implementation and operation:”data awareness” that matters. Annals of translational medicine, 8(7):501, 2020.

Dratsch, T., Chen, X., Rezazade Mehrizi, M., Kloeckner, R., Mähringer-Kunz, A., Püsken, M., Baeßler, B., Sauer, S., Maintz, D., and Pinto dos Santos, D. Automation bias in mammography: the impact of artificial intelligence bi-rads suggestions on reader performance. Radiology, 307(4):e222176, 2023.

Fischl, B. Freesurfer. Neuroimage, 62(2):774–781, 2012.

Gorgolewski, K. J., Alfaro-Almagro, F., Auer, T., Bellec, P., Capotă, M., Chakravarty, M. M., Churchill, N. W., Cohen, A. L., Craddock, R. C., Devenyi, G. A., et al. Bids apps: Improving ease of use, accessibility, and reproducibility of neuroimaging data analysis methods. PLoS computational biology, 13 (3):e1005209, 2017.

Isensee, F., Schell, M., Pflueger, I., Brugnara, G., Bonekamp, D., Neuberger, U., Wick, A., Schlemmer, H.-P., Heiland, S., Wick, W., et al. Automated brain extraction of multisequence mri using artificial neural networks. Human brain mapping, 40(17):4952–4964, 2019.

Isensee, F., Jaeger, P. F., Kohl, S. A., Petersen, J., and Maier-Hein, K. H. nnu-net: a self-configuring method for deep learning-based biomedical image segmentation. Nature methods, 18(2):203–211, 2021.

Kelly, C. J., Karthikesalingam, A., Suleyman, M., Corrado, G., and King, D. Key challenges for delivering clinical impact with artificial intelligence. BMC medicine, 17(1):195, 2019.

Liu, J., Xia, C. S., Wang, Y., and Zhang, L. Is your code generated by chatgpt really correct? rigorous evaluation of large language models for code generation. Advances in neural information processing systems, 36:21558–21572, 2023.

Lu, M. Y., Williamson, D. F., Chen, T. Y., Chen, R. J., Barbieri, M., and Mahmood, F. Data-efficient and weakly supervised computational pathology on whole-slide images. Nature biomedical engineering, 5(6):555–570, 2021.

Moassefi, M., Rouzrokh, P., Conte, G. M., Vahdati, S., Fu, T., Tahmasebi, A., Younis, M., Farahani, K., Gentili, A., Kline, T., et al. Reproducibility of deep learning algorithms developed for medical imaging analysis: a systematic review. Journal of digital imaging, 36(5):2306–2312, 2023.

Muehlematter, U. J., Daniore, P., and Vokinger, K. N. Approval of artificial intelligence and machine learning-based medical devices in the usa and europe (2015–20): a comparative analysis. The Lancet Digital Health, 3(3):e195–e203, 2021.

Ouyang, S., Zhang, J. M., Harman, M., and Wang, M. An empirical study of the non-determinism of chatgpt in code generation. ACM Transactions on Software Engineering and Methodology, 34(2):1–28, 2025.

Patil, S. G., Zhang, T., Wang, X., and Gonzalez, J. E. Gorilla: Large language model connected with massive apis. Advances in Neural Information Processing Systems, 37:126544–126565, 2024.

Pimentel, J. F., Murta, L., Braganholo, V., and Freire, J. A large-scale study about quality and reproducibility of jupyter notebooks. In 2019 IEEE/ACM 16th international conference on mining software repositories (MSR), pp. 507–517. IEEE, 2019.

Qin, Y., Liang, S., Ye, Y., Zhu, K., Yan, L., Lu, Y., Lin, Y., Cong, X., Tang, X., Qian, B., et al. Toolllm: Facilitating large language models to master 16000+ real-world apis. In International Conference on Learning Representations, volume 2024, pp. 9695–9717, 2024.

Schick, T., Dwivedi-Yu, J., Dessì, R., Raileanu, R., Lomeli, M., Hambro, E., Zettlemoyer, L., Cancedda, N., and Scialom, T. Toolformer: Language models can teach themselves to use tools. Advances in neural information processing systems, 36:68539–68551, 2023.

Sclar, M., Choi, Y., Tsvetkov, Y., and Suhr, A. Quantifying language models’ sensitivity to spurious features in prompt design or: How i learned to start worrying about prompt formatting. In International Conference on Learning Representations, volume 2024, pp. 25055–25083, 2024.

Simkó, A., Garpebring, A., Jonsson, J., Nyholm, T., and Löfstedt, T. Reproducibility of the methods in medical imaging with deep learning. In Medical Imaging with Deep Learning, pp. 95–106. PMLR, 2024.

Spracklen, J., Wijewickrama, R., Sakib, A. N., Maiti, A., and Viswanath, B. We have a package for you! a comprehensive analysis of package hallucinations by code generating {LLMs}. In 34th USENIX Security Symposium (USENIX Security 25), pp. 3687–3706, 2025.

Tanno, R., Barrett, D. G., Sellergren, A., Ghaisas, S., Dathathri, S., See, A., Welbl, J., Lau, C., Tu, T., Azizi, S., et al. Collaboration between clinicians and vision–language models in radiology report generation. Nature Medicine, 31(2):599–608, 2025.

Tian, Y., Yan, W., Yang, Q., Zhao, X., Chen, Q., Wang, W., Luo, Z., Ma, L., and Song, D. Code-halu: Investigating code hallucinations in llms via execution-based verification. In Proceedings of the AAAI Conference on Artificial Intelligence, volume 39, pp. 25300–25308, 2025.

Trisovic, A., Lau, M. K., Pasquier, T., and Crosas, M. A large-scale study on research code quality and execution. Scientific Data, 9(1):60, 2022.

Tustison, N. J., Cook, P. A., Holbrook, A. J., Johnson, H. J., Muschelli, J., Devenyi, G. A., Duda, J. T., Das, S. R., Cullen, N. C., Gillen, D. L., et al. The antsx ecosystem for quantitative biological and medical imaging. Scientific reports, 11(1):9068, 2021.

Varoquaux, G. and Cheplygina, V. Machine learning for medical imaging: methodological failures and recommendations for the future. NPJ digital medicine, 5(1):48, 2022.

Von Chamier, L., Laine, R. F., Jukkala, J., Spahn, C., Krentzel, D., Nehme, E., Lerche, M., Hernández-Pérez, S., Mattila, P. K., Karinou, E., et al. Democratising deep learning for microscopy with zerocostdl4mic. Nature communications, 12(1):2276, 2021.

Wasserthal, J., Breit, H.-C., Meyer, M. T., Pradella, M., Hinck, D., Sauter, A. W., Heye, T., Boll, D. T., Cyriac, J., Yang, S., et al. Totalsegmentator: robust segmentation of 104 anatomic structures in ct images. Radiology: Artificial Intelligence, 5(5): e230024, 2023.

Yao, S., Zhao, J., Yu, D., Du, N., Shafran, I., Narasimhan, K., and Cao, Y. React: Synergizing reasoning and acting in language models. In International Conference on Learning Representations, volume 2023, 2023.

Zhang, Z., Wang, C., Wang, Y., Shi, E., Ma, Y., Zhong, W., Chen, J., Mao, M., and Zheng, Z. Llm hallucinations in practical code generation: Phenomena, mechanism, and mitigation. Proceedings of the ACM on Software Engineering, 2(ISSTA):481–503, 2025.

